# Respiratory dysfunctions in two rodent models of chronic epilepsy and acute seizures and their link with brainstem serotonin system

**DOI:** 10.1101/2021.04.11.439321

**Authors:** Hayet Kouchi, Michaël Ogier, Gabriel Dieuset, Anne Morales, Béatrice Georges, Jean-Louis Rouanet, Benoît Martin, Philippe Ryvlin, Sylvain Rheims, Laurent Bezin

## Abstract

Patients with drug-resistant epilepsy can experience respiratory alterations notably during seizures. The mechanisms underlying this long-term alteration of respiratory function remain unclear. This study aimed at determining in rats whether epilepsy is associated with alterations of both the respiratory function and brainstem serotonin (5-HT) system. Epilepsy was triggered by pilocarpine-induced *status epilepticus* in rats. 30-50% of epileptic (EPI) rats exhibited sharp decrease of oxygen consumption (SDOC), low metabolic rate of oxygen and slow regular ventilation; these rats were called EPI/SDOC+ rats. These alterations were only detected in rats with chronic epilepsy, independent of behavioral seizures, persisted over the time, and were not associated with death. In these rats, 5-HT fiber density in the nucleus *tractus solitarius* was below that of control and EPI/SDOC-rats. Both EPI/SDOC+ rats and DBA/2 mice presenting with fatal respiratory arrest following an audiogenic-induced seizure, a model of sudden and expected death in epilepsy, had increased transcript levels of tryptophan hydroxylase 2 (p<0.001 for both strains) and 5-HT presynaptic transporter (rats: p=0.003; mice: p=0.001). Thus, our data support that 5-HT alterations are associated with chronic and acute epilepsy-related respiratory dysfunctions.

## Introduction

Seizure-related respiratory dysfunctions in patients with epilepsy are well characterized and can be observed either during focal seizures or during generalized tonic-clonic seizures (GTCS) [1, 2]. In patients with drug-resistant epilepsy, oxygen desaturation <90%, as measured by pulse oximetry, has thus been reported in 33% of focal seizures [3] and in up to 86% of GTCS [2]. In both seizure types, these transient hypoxemias result from ictal and/or post-ictal central apnea [1, 4]. In addition, respiratory dysfunctions are not only limited to seizure periods in some patients with epilepsy. Obstructive sleep apnea syndrome (OSA) is thus prevalent in people with drug-resistant epilepsy ranging between 30-49% and markedly exceeding the estimated rate in the general population [5, 6].

Importantly, these respiratory dysfunctions might represent a risk factor for sudden unexpected death in epilepsy (SUDEP), which primarily results from post-ictal central apnea [3], urging to fill the gap of knowledge regarding epilepsy-related respiratory alteration and its pathological mechanisms. Post-mortem examination of the brainstem of patients who died from SUDEP showed changes in the ventrolateral medulla (VLM) [7], a region containing different populations of respiratory neurons including those of the pre-Bötzinger Complex, a kernel generator of the inspiratory rhythm [8]. Specifically, SUDEP victims presented with an alteration of brainstem serotonin (5-HT) system with notable decrease in both tryptophan hydroxylase (TPH2), the rate-limiting enzyme in 5-HT synthesis, and 5-HT presynaptic transporter (SERT) within medullary raphe nuclei and/or the VLM [7].

Similarly, brainstem 5-HT abnormalities have been reported in DBA/1 and DBA/2 mouse strains, which are frequently used as a SUDEP model in preclinical studies because they can exhibit a fatal respiratory arrest (FRA) after induction of an audiogenic seizure (AGS) [9–11].

Interestingly, FRAs are prevented when DBA/2 and DBA/1 mice are treated with fluoxetine, a selective serotonin reuptake inhibitor (SSRI) before AGS induction, which is consistent with the hypothesis that 5-HT system may have a strong implication in FRA following AGS [12, 13]. Nevertheless, animals prone to AGS-induced FRA present some limitations to completely characterize respiratory dysfunction in chronic epilepsy and its potential associated 5-HT abnormalities, mostly because they represent models of induced seizure(s) and do not develop spontaneous recurrent seizures.

In this study, we used one of the most studied rat model of chronic epilepsy, following pilocarpine-induced *status epilepticus* (Pilo-SE). As for a large number of patients with temporal lobe epilepsy, Pilo-SE rats are notably characterized by chronic, spontaneous and drug-resistant seizures [14]. In this model, while mortality is present during SE, no mortality has been reported once chronic phase of epilepsy has developed. Thus, to further explore the hypothesis that epilepsy-related respiratory alterations are associated with altered brainstem 5-HT system, we first explored whether respiratory alterations occurred with development of epilepsy, and then investigated whether they were associated with brainstem 5-HT system alterations; i.e. number of 5-HT-positive cell bodies, density of 5-HT-positive terminals, TPH2 and SERT mRNA levels. To help better conclude on the potential involvement of 5-HT system alterations with respect to the respiratory disorders observed during the chronic phase of epilepsy and independently of any risk of SUDEP-like mortality, we finally compared brainstem TPH2 and SERT mRNA levels between EPI rats and DBA/2 mice exhibiting FRA after AGS.

## Results

### Development of behavioral spontaneous recurrent seizures (SRSs) in rats after Pilo-SE

The numbers of rats that did not develop Pilo-SE or died during SE are reported in the Supplementary information section. SRSs were identified as detailed in the Methods section. Briefly, out of the 82 rats subjected to Pilo-SE in experiments 1-3, 59 developed at least two consecutive SRSs by the end of the 2^nd^ week post-SE, and were then stated as EPI rats.

According to the evaluation of two experimenters, the number of recorded seizures (stages 3-5 of severity according to Racine’s scale) per 12 hours during the daylight period was about 1.1 ± 0.1 in EPI rats from weeks 3 to 8 after SE. In experiment 1, none of the Pilo-SE rats had developed 2 consecutive Handling-Induced Seizures (HIS) at one-week post-SE, in agreement with previous studies [15].

### A subset of EPI rats presented with abnormal pattern of oxygen consumption

In experiment 1, metabolic rate of oxygen 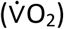 of 20 EPI rats and 8 controls was recorded for 24h each week from weeks 1 to 8 after SE. A thorough screening of 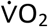 was then made for each rat.

#### Presence of sharp decreases in oxygen consumption (SDOCs)

Control rats (n=8) presented with slight up and down fluctuations of 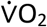 during both light and dark periods, as illustrated in one rat (Figure 1A). For a better characterization of 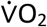 fluctuations within the daily pattern, 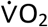 was averaged for each rat during the light and dark periods, separately. Afterwards, the lowest value below the averaged 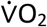 was determined for each rat; it ranged between 38% and 70% of their average 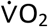, independently of the circadian rhythm. This pattern remained unchanged during the next 7 weeks of recordings. In rats subjected to Pilo-SE, the pattern of daily 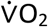 was similar to that of controls during the first two weeks post-SE, as illustrated in one Pilo-SE rat (Figure 1B). However, by the end of the 2^nd^ week post-SE, two groups of EPI rats could be identified according to 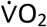 fluctuations. A subset of EPI rats presented with a large amplitude of 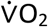 decrease (“Sharp Decrease(s) in Oxygen Consumption”; SDOCs), defined as a decrease greater than −62%, a percentage that corresponds to the maximal decrease of 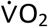 in control rats. Such EPI rats were thus identified as EPI/SDOC+ rats and their lowest 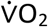 values ranged between 0% and 32% of the average 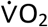, as illustrated at 3 weeks post-SE in one rat (Figure 1C). By contrast, the remaining EPI rats (EPI/SDOC-rats) were similar to controls with regard to the lowest 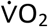 values, which ranged between 41% and 67% of the average 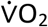. Video monitoring confirmed that SDOCs were not associated with behavioral seizures, as illustrating while rat adopted a sleep-like position (Figure 2A). By contrast, behavioral seizures were associated with a long-lasting increase of the 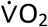, as illustrated during a stage-5 seizure (Figure 2B).

**Figure 1:**
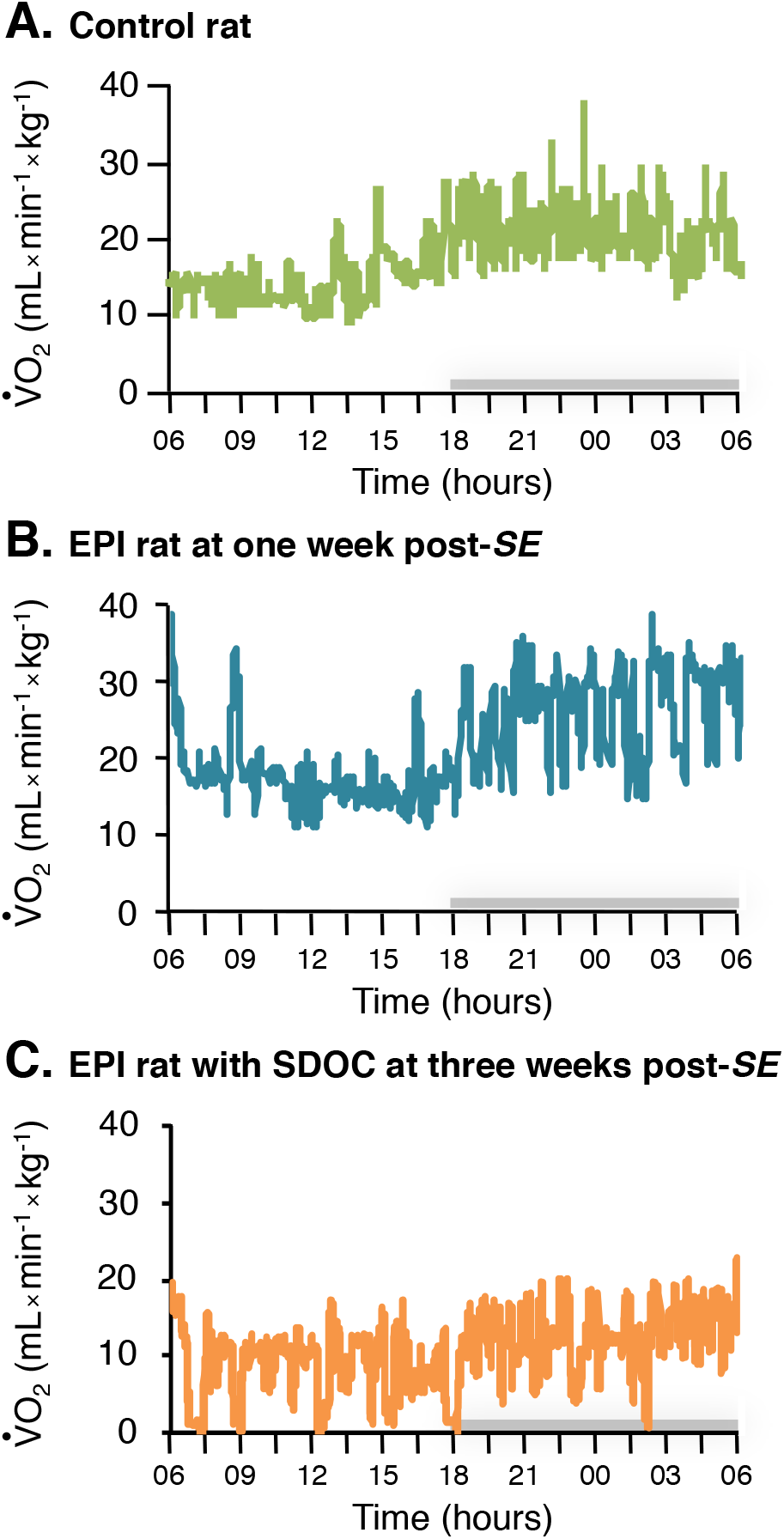
Detection of SDOC events during a 24-hour monitoring of oxygen consumption in a subgroup of epileptic rats. Oxygen consumption 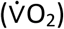 (light period from 06:00 am to 6:00 pm) was measured using respiratory thermochemistry. Grey bar above each x-axis indicates the dark period. A single profile was observed in control rats (A), while two distinct profiles were distinguished in epileptic rats (B, C). The first profile was similar to that observed in control rats (B), the second one was characterized by several episodes of sharp decrease in oxygen consumption (SDOC) (C), sometimes reaching the zero value. These SDOC events were only seen in epileptic rats, thereby discriminating two groups of epileptic rats upon their presence (EPI/SDOC+) or their absence (EPI/SDOC-).

**Figure 2:**
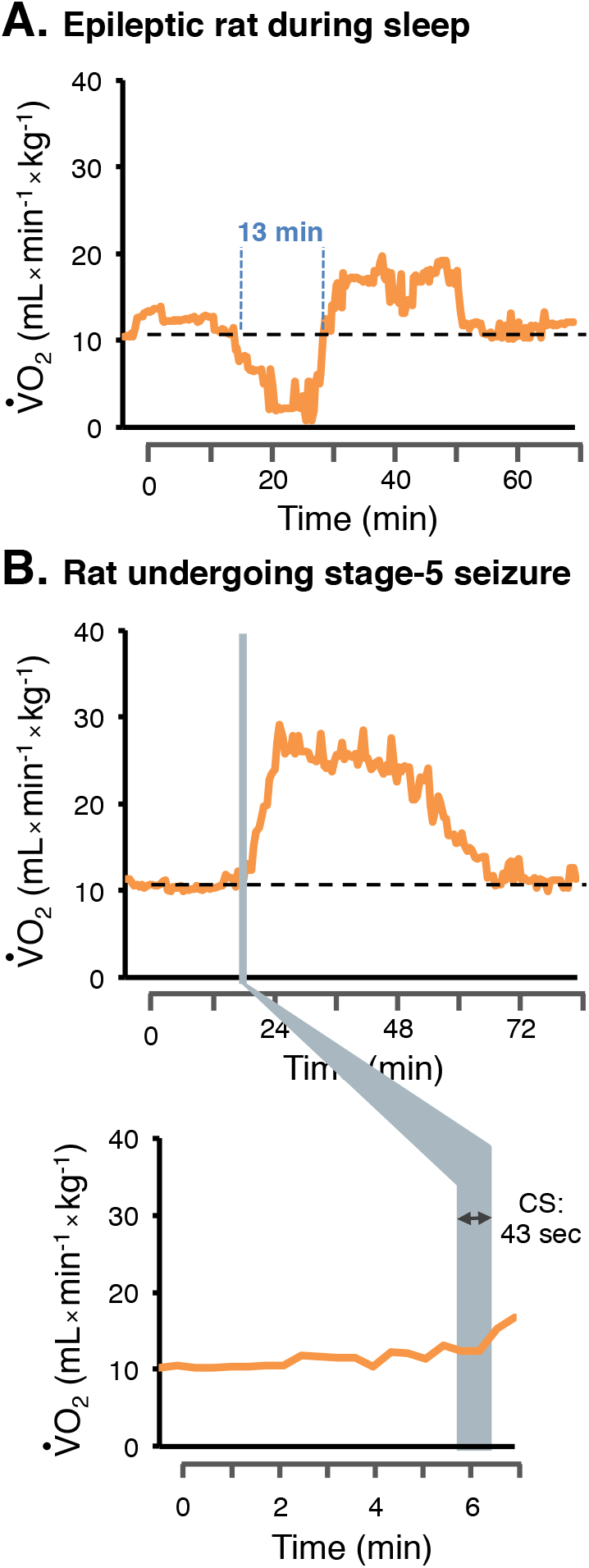
SDOC can occur independently of behavioral spontaneous seizures. (A) Example of a SDOC event recorded in an EPI rat at 5 weeks post-SE, when it was adopting a sleep posture. This SDOC event occurred during the light period and was identified with the decline in 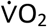 values. No behavioral seizure was observed before, during and after the SDOC occurrence. By contrast, (B) a sharp increase in 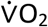 values was observed after a 43-second stage-5 seizure according to Racine’s scale in an EPI/SDOC+ rat whose values of 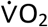 were above pre-ictal level for about 30 minutes.

Regarding the highest 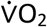 values in control rats, they were ranged between 164% and 229% of the average 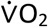 (median=183%). The highest fluctuations within the daily 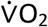 pattern of EPI/SDOC+ and EPI/SDOC-rats were not statistically different from controls, and ranged between 144% and 227% of the average 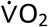 (median=180%). However, EPI/SDOC+ rats had greater increases of 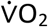 than those observed in EPI/SDOC-rats (+93 ± 5% *vs* +75 ± 4%; P=0.01).

As a good indicator of the stability of 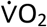 pattern, the intra-individual variation (coefficient of variation, CV) of the daily oxygen consumption pattern (CV-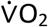) was calculated independently of light and dark periods. The mean CV-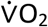 of the control group was estimated to be 20.8 ± 0.5% and 22 ± 0.7% for the light and the dark periods, respectively. The mean CV-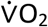 of EPI/SDOC-rats was almost similar to that of controls with 19.1 ± 0.8% and 22.8 ± 0.7% for the light and dark periods, respectively. However, in EPI/SDOC+ rats, it was significantly higher than that measured in both control (P= 0.002 and p= 0.004 for light and dark periods, respectively) and EPI/SDOC-groups (P< 0.0001 and p= 0.017 for light and dark periods, respectively), reaching 27.4 ± 2.9% and 26.5 ± 1.3% for light and dark periods, respectively.

#### Low metabolic rate of oxygen in EPI/SDOC+ rats

Since 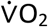 was measured weekly for all rats for 8 weeks, it was possible to retrospectively assign each EPI rat into either the EPI/SDOC+ or the EPI/SDOC-group from the first week after SE.

We investigated the evolution of the daily 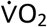 in each group. We first verified that SDOC events in EPI/SDOC+ rats did not affect the average value of the daily 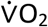, both during the light (P=0.86) and the dark (P=0.93) periods. Indeed, in order to compare between the three groups of rats the daily 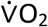 during the 8 weeks of measurement, we preferred to use the data without SDOCs in EPI/SDOC+ rats. We show in control and EPI/SDOC-rat groups that the daily 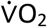 decreased progressively by 21.3 ± 2.4% (P=0.05) and 22.0 ± 2.2% (P=0.02), respectively, between weeks 1-2 and 8 post-SE. By contrast, the decrease observed in EPI/SDOC+ rats was much larger during the same period (−69.5 ± 1.1%; P=0.002), and already reached −53 ± 6.0% (P=0.02) by the fifth week post-SE (Figure 3A). We also found that the decrease of daily 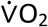 coincided with the onset of SDOCs (Figure 3B).

**Figure 3:**
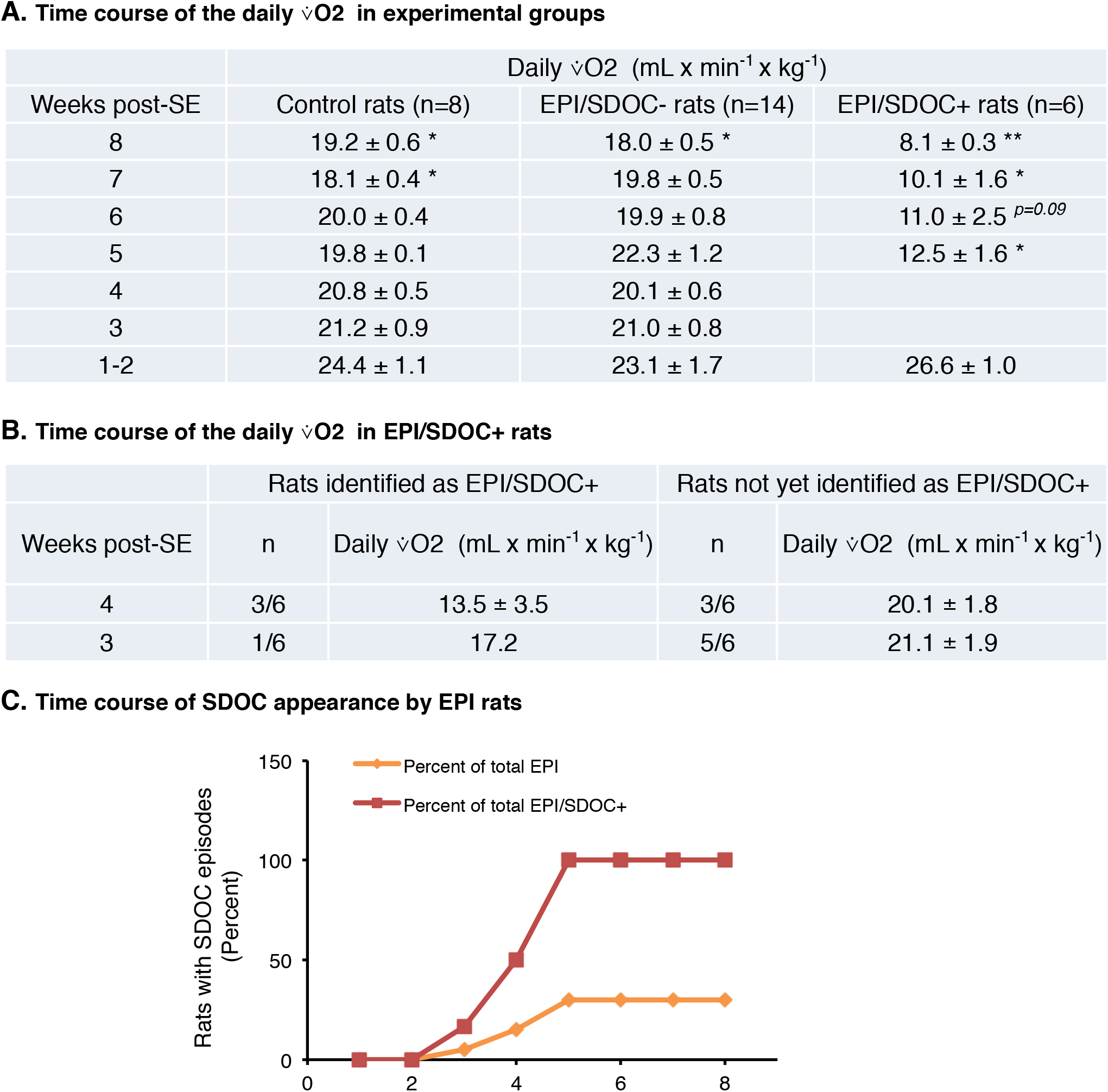
Time course of daily 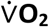 pattern. Respiratory thermochemistry was recorded once a week for each rat, from the 1^st^ to the 8^th^ week post-SE. (A) Daily 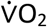 is calculated for each group (control rats (n=8), EPI/SDOC-rats (n=14), EPI/SDOC+ rats (n=6)) from the 1 to 8 weeks post-SE. (B) The daily 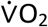 of animals is calculated at 3-4 weeks post-SE in animals identified or not yet as EPI/SDOC+ rats. Paired comparison of daily 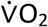 over weeks showed statistical significance with only 1-2 weeks post-SE. Results are expressed in mean ± SEM of 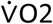, *, p<0.05; **, P=0.01 in comparison of daily 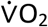 at 1-2 weeks post-SE. (C) the first SDOC was detected in one rat (1/20) during the 3rd post-SE week. The maximum number of EPI rats exhibiting SDOCs (EPI/SDOC+, 30%) was reached by the end of the 5^th^ week post-SE. The proportion of EPI/SDOC+ rats was expressed in percent of the total number of rats developing epilepsy (yellow curve) or in percent of the total number of rats presenting with SDOCs by the end of the 8^th^ post-SE (red curve).

In each group, we verified that the daily 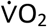 did not change significantly between 5 and 8 weeks post-SE. Then, the daily 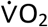 has been averaged during that period for each group to ease the comparison. We found that the daily 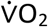 was greatly reduced in EPI/SDOC+ rats at 5-8 weeks post-SE compared to both EPI/SDOC-rats and controls (−39 ± 7 % and −34 ± 7 %, respectively). This decrease was observed during both the light period (EPI/SDOC+ *vs*. EPI/SDOC-rats: p< 0.001; EPI/SDOC+ *vs*. control rats: P=0.002) and the dark period (EPI/SDOC+ *vs*. EPI/SDOC-rats: p< 0.001; EPI/SDOC+ *vs*. control rats: P=0.002) (Supplementary Figure S1A). At weeks 1-2 post-SE, no difference was observed between groups during both light and dark periods (Supplementary Figure S1A).

In the 3 groups of rats (controls, EPI/SDOC+ and EPI/SDOC-), we evaluated the effect of three factors that have been referenced as affecting 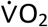 [16]: time (t_0_ corresponds to the time of induction of SE in EPI rats), body weight and circadian rhythm. Full description of these effects is detailed in the Supplementary information section.

#### Characterization of SDOCs

The number of SDOCs during the 24-hour measurement ranged between 12 and 414 events (median = 44), with no difference of SDOC numbers between dark and light periods. The median duration of SDOCs was 5 min (range: 20 sec to 13 min). The longest SDOC event is illustrated in figure 2A; it lasted 13 min and occurred, according to the video, during the daylight period while the animal was sleeping without any evidence of behavioral seizure. Full description of 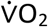 pattern during the manifestation of seizures is detailed in the Supplementary information section.

Body position during SDOCs was far different from freezing posture frequently observed before rats had a behavioral seizure that was easily discernible by visual observation alone (stages 3 to 5 on Racine’s scale). It is also noteworthy that rats always exhibited a quiet state during SDOC events. The proportion of EPI/SDOC+ rats increased between weeks 3 and 5 post-SE and then stabilized (from weeks 5 to 8 post-SE). By the end of the 5^th^ week post-SE, 30% (6/20) of EPI rats exhibited SDOCs (Figure 3C).

### Ventilatory and cardiac functions in epileptic rats

Experiment 2 aimed to further investigate respiratory function by evaluating ventilatory and cardiac functions of EPI rats during chronic phase of epilepsy. It should be noted that we could not record respiratory variables using whole-body plethysmography before the 10^th^ week post-SE because rats were not able to stay calm in the apparatus chamber, likely due to enhanced reactivity to stressful situations.

#### EPI rats presented with a low steady respiratory pattern and enhanced ventilatory response to hypoxia and hypercapnia

Under normoxia, EPI rats (n=16) presented a slow respiratory pattern with a significant decrease of minute ventilation 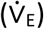 compared to controls (n=5) (−12 ± 3%, P=0.03) (Supplementary Figure S2A). The reduction of 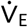 in EPI rats resulted from a decrease of respiratory frequency (f_R_) (−12 ± 2% *vs* controls, P=0.001) while tidal volume (V_T_) was similar to controls (P=0.92). The decrease of f_R_ of EPI rats was related to a simultaneous increase of inspiratory time (T_I_) and expiratory time (T_E_), that reached +20 ± 3% (P=0.01) and +10 ± 2 % (P=0.001) compared to the values measured in controls, respectively. Consequently, the total respiratory cycle duration (T_tot_) of EPI rats was significantly higher than controls (+14 ± 2 %; P=0.002) (Supplementary Figure S2A).

Regarding the stability of breathing pattern, the coefficient of variation of breathing, apnea frequency and duration were similar between EPI and control rats (p=0.16, p=0.61 and p=0.09, respectively) (Supplementary Figure S2B).

Compared to control rats (n=13), EPI rats (n=18) showed a greater hypoxia ventilatory response (HVR) with higher 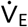 (+23 ± 6 % *vs* controls, P=0.007), higher f_R_ (+16 ± 2 % *vs* controls, P<0.001) and similar V_T_ (P=0.06). Similarly, the hypercapnia ventilatory response (HCVR) of EPI rats (n=18) was higher than controls (n=12) with greater 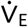 (+14 ± 4 % *vs* controls, P=0.01) and greater f_R_ (+10 ± 3 % *vs* controls, p=0.02), while V_T_ remained similar to that of controls (P=0.09).

#### Abnormal 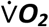 pattern is associated with pronounced decrease of ventilatory rhythm at rest

Since oxygen is strictly provided during ventilatory process, we aimed to compare ventilation between EPI rats with or without abnormal 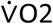 pattern. In experiment 2, we were able to discriminate 8 EPI/SDOC+ rats, which presented with a daily 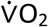 lower than that of both EPI/SDOC-rats (n=15, −51 ± 9%; P< 0.001) and control rats (n=16, −41 ± 11%; P=0.003).

Under normoxia, the ventilation of EPI/SDOC-rats was significantly altered compared to control rats (Table 1A), but to a lesser extent than that of EPI/SDOC+ rats. Indeed, compared to control rats, EPI/SDOC-rats presented with a slight but not significant 6 ± 3% reduction of 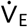, associated with an 8 ± 2% reduction (P=0.015) of f_R_, a 13 ± 1% increase (P<0.001) of T_I_ and a 9 ± 2% increase (P=0.025) of T_tot_ (Table 1A). Furthermore, compared to EPI/SDOC-rats, EPI/SDOC+ rats presented with an additional 13 ± 3% reduction (P=0.013) of 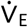, that resulted from an 8 ± 2% reduction (P=0.036) of f_R_, a 13 ± 3% increase (P<0.001) of T_I_ and an 8 ± 3% increase (P=0.013) of T_tot_ (Table 1A). Tidal volume (V_T_), the coefficient of variation of the breathing frequency, apnea frequency and duration were similar in all groups (Table 1A). Under hypoxic and hypercapnic conditions, 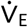, V_T_, and f_R_ was similar between EPI/SDOC+ (n=6) and EPI/SDOC-rats (n=12) (Table 1B).

**Table 1:**
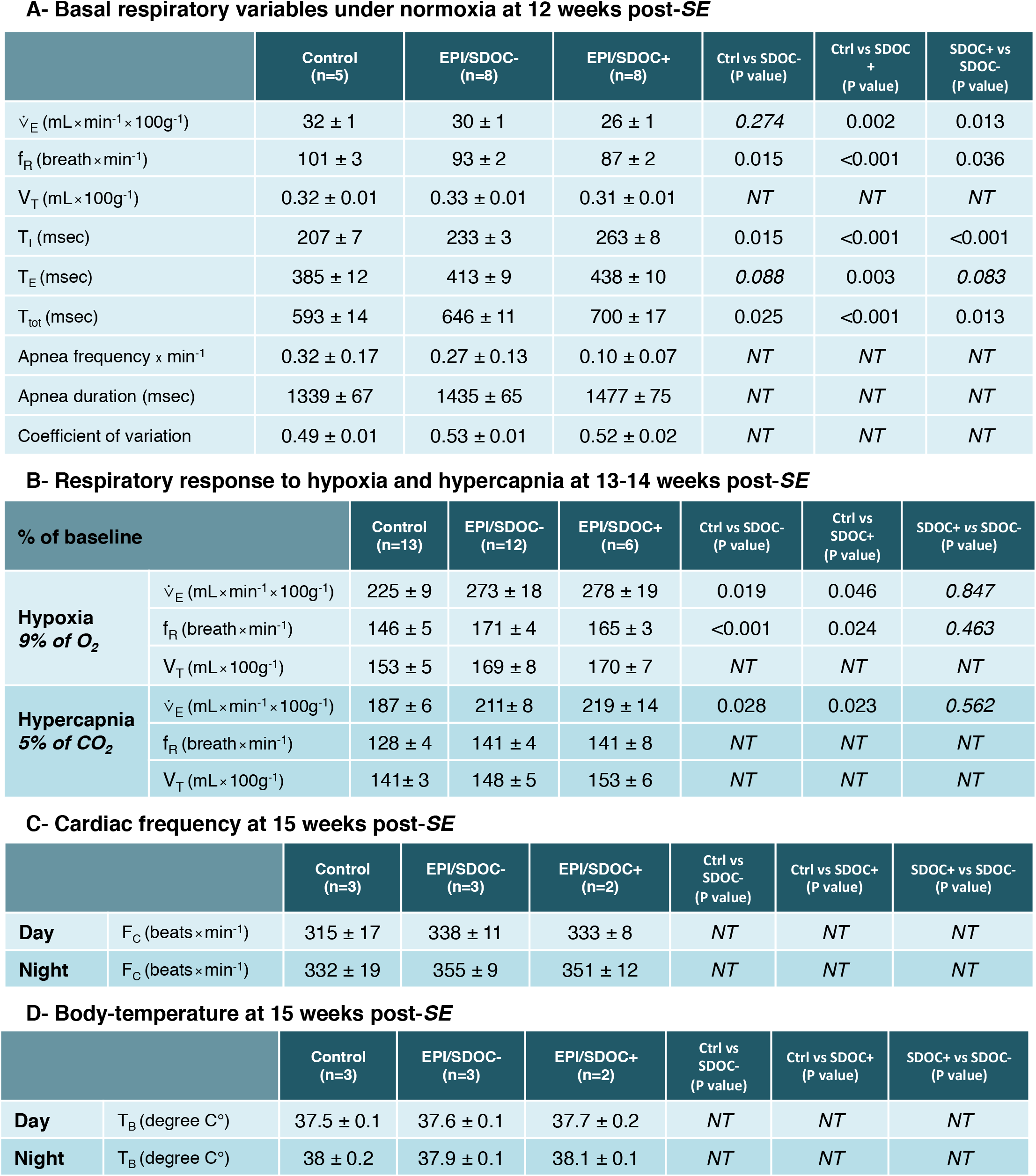
Monitoring of cardio-respiratory functions in control and EPI rats between 12 and 15 weeks post-SE. Using plethysmography, the ventilatory variables were measured under normoxic condition (21% O_2_) at 12 weeks post-SE (A) and in response to hypoxia (9% O_2_) or hypercapnia (5% CO_2_) exposures between 13-14 weeks post-SE (B). Cardiac frequency (C) and body-temperature (D) were concomitantly evaluated at 15 weeks post-SE. Abbreviations: Fc, cardiac frequency; f_R_, respiratory frequency; T_B_, body-temperature; T_E_, expiration time; T_I_, inspiration time; T_tot_, total respiratory cycle duration; 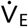, minute volume; V_T_, tidal volume. Statistics: NT, not statistically significant.

A- Basal respiratory variables under normoxia at 12 weeks post-SE
|  | Control<br>(n=5) | EPI/SDOC-<br>(n=8) | EPI/SDOC+<br>(n=8) | Ctrl vs SDOC-<br>(P value) | Ctrl vs SDOC+<br>(P value) | SDOC+ vs SDOC-<br>(P value) |
| --- | --- | --- | --- | --- | --- | --- |
| $\dot{V}_E$ (mL×min$^{-1}\times$100g$^{-1}$) | 32 ± 1 | 30 ± 1 | 26 ± 1 | 0.274 | 0.002 | 0.013 |
| $f_R$ (breath×min$^{-1}$) | 101 ± 3 | 93 ± 2 | 87 ± 2 | 0.015 | <0.001 | 0.036 |
| $V_T$ (mL×100g$^{-1}$) | 0.32 ± 0.01 | 0.33 ± 0.01 | 0.31 ± 0.01 | NT | NT | NT |
| $T_I$ (msec) | 207 ± 7 | 233 ± 3 | 263 ± 8 | 0.015 | <0.001 | <0.001 |
| $T_E$ (msec) | 385 ± 12 | 413 ± 9 | 438 ± 10 | 0.088 | 0.003 | 0.083 |
| $T_{tot}$ (msec) | 593 ± 14 | 646 ± 11 | 700 ± 17 | 0.025 | <0.001 | 0.013 |
| Apnea frequency × min$^{-1}$ | 0.32 ± 0.17 | 0.27 ± 0.13 | 0.10 ± 0.07 | NT | NT | NT |
| Apnea duration (msec) | 1339 ± 67 | 1435 ± 65 | 1477 ± 75 | NT | NT | NT |
| Coefficient of variation | 0.49 ± 0.01 | 0.53 ± 0.01 | 0.52 ± 0.02 | NT | NT | NT |

B- Respiratory response to hypoxia and hypercapnia at 13-14 weeks post-SE
| % of baseline |  | Control<br>(n=13) | EPI/SDOC-<br>(n=12) | EPI/SDOC+<br>(n=6) | Ctrl vs SDOC-<br>(P value) | Ctrl vs SDOC+<br>(P value) | SDOC+ vs SDOC-<br>(P value) |
| --- | --- | --- | --- | --- | --- | --- | --- |
| **Hypoxia**<br>9% of O<sub>2</sub> | $\dot{V}_E$ (mL×min$^{-1}\times$100g$^{-1}$) | 225 ± 9 | 273 ± 18 | 278 ± 19 | 0.019 | 0.046 | 0.847 |
| | $f_R$ (breath×min$^{-1}$) | 146 ± 5 | 171 ± 4 | 165 ± 3 | <0.001 | 0.024 | 0.463 |
| | $V_T$ (mL×100g$^{-1}$) | 153 ± 5 | 169 ± 8 | 170 ± 7 | NT | NT | NT |
| **Hypercapnia**<br>5% of CO<sub>2</sub> | $\dot{V}_E$ (mL×min$^{-1}\times$100g$^{-1}$) | 187 ± 6 | 211 ± 8 | 219 ± 14 | 0.028 | 0.023 | 0.562 |
| | $f_R$ (breath×min$^{-1}$) | 128 ± 4 | 141 ± 4 | 141 ± 8 | NT | NT | NT |
| | $V_T$ (mL×100g$^{-1}$) | 141 ± 3 | 148 ± 5 | 153 ± 6 | NT | NT | NT |

C- Cardiac frequency at 15 weeks post-SE
|  |  | Control<br>(n=3) | EPI/SDOC-<br>(n=3) | EPI/SDOC+<br>(n=2) | Ctrl vs SDOC-<br>(P value) | Ctrl vs SDOC+<br>(P value) | SDOC+ vs SDOC-<br>(P value) |
| --- | --- | --- | --- | --- | --- | --- | --- |
| **Day** | $F_C$ (beats×min$^{-1}$) | 315 ± 17 | 338 ± 11 | 333 ± 8 | NT | NT | NT |
| **Night** | $F_C$ (beats×min$^{-1}$) | 332 ± 19 | 355 ± 9 | 351 ± 12 | NT | NT | NT |

**Table 1:** Monitoring of cardio-respiratory functions in control and EPI rats between 12 and 15 weeks post-SE.
|  |  | Control<br>(n=3) | EPI/SDOC-<br>(n=3) | EPI/SDOC+<br>(n=2) | Ctrl vs SDOC-<br>(P value) | Ctrl vs SDOC+<br>(P value) | SDOC+ vs SDOC-<br>(P value) |
| --- | --- | --- | --- | --- | --- | --- | --- |
| **Day** | $T_B$ (degree C$^{\circ}$) | 37.5 ± 0.1 | 37.6 ± 0.1 | 37.7 ± 0.2 | NT | NT | NT |
| **Night** | $T_B$ (degree C$^{\circ}$) | 38 ± 0.2 | 37.9 ± 0.1 | 38.1 ± 0.1 | NT | NT | NT |

#### No alteration of cardiac frequency and body-temperature in EPI rats

Respiratory system, cardiac function and body-temperature have a tight relationship. In this experiment, we showed that cardiac frequency and body-temperature of EPI/SDOC+ rats (n=2), EPI/SDOC-rats (n=3) and control rats (n=3) were similar during both light and dark periods (Table 1C, 1D).

### Brainstem 5-HT system of EPI rats

In experiment 3, 20 rats developed epilepsy. Among those rats, we identified 8 EPI/SDOC+ and 8 EPI/SDOC-rats. Eight control rats were used. Brainstem was processed for either histological (n=3/8 rats in each group) or gene expression (n=5/8 rats in each group) analyses.

#### Alteration of the density of brainstem 5-HT immunopositive processes in EPI rats

We determined whether metabolic and ventilatory alterations developed by EPI/SDOC+ rats were associated with dysfunction of the 5-HT system in the brainstem. Therefore, we focused our investigation within caudal raphe nuclei, which are known to innervate neuronal groups closely implicated in the regulation of respiratory rhythm generation, such as the NTS and the VLM. 5-HT-immunolabeling was detected in both processes and cell bodies. Within caudal raphe nuclei (pallidus, magnus and obscurus), although we observed a downward trend in epileptic rats, there was no statistical difference in the number of 5-HT-positive neuronal cell bodies between the three groups of rats (control rats: 22 ± 6, EPI/SDOC-: 18 ± 4, EPI/SDOC+: 13 ± 5, P=0.339). In addition, there was no difference in surface area of 5-HT positive cell bodies (control rats: 383 ± 21 pixel^2^, EPI/SDOC-rats: 445 ± 37 pixel^2^, EPI/SDOC+ rats: 397 ± 54 pixel^2^, P=0.383) nor in the index of 5-HT neuronal concentration (control rats: 109 ± 19 A.U., EPI/SDOC-rats: 125 ± 23 A.U., EPI/SDOC+ rats: 119 ± 18 A.U., P=0.864) between the three rat groups. The density of 5-HT immunopositive processes within both the VLM (Figure 4A) and the NTS (Figure 4B) was significantly higher in EPI/SDOC-rats compared to EPI/SDOC+ rats and control rats (Figures 4A4 and 4B4). Furthermore, the density of 5-HT-immunolabeled processes within the NTS was significantly lower in EPI/SDOC+ rats compared to controls (Figure 4B4).

**Figure 4:**
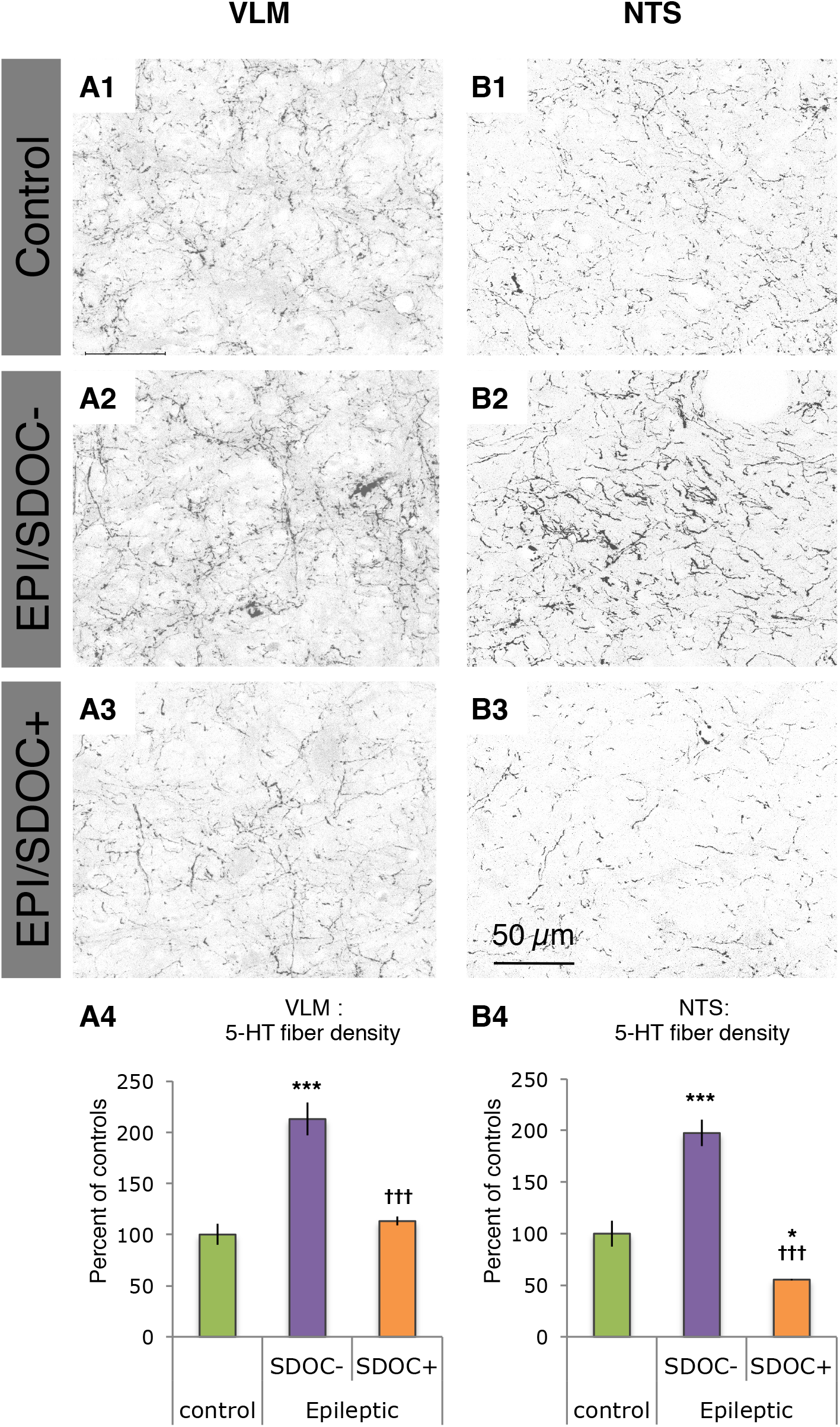
Semi-quantification of 5-HT immunopositive terminals of caudal raphe nuclei within the medulla oblongata of control and EPI rats. Fluorescent 5-HT labeling was performed on sections selected at interaural −2.60 mm within two brainstem regions involved in the regulation of respiratory rhythm generation: the ventrolateral medulla (VLM) and nucleus tractus solitarius (NTS). Images of 5-HT positive fibers were captured within the VLM (A1, A2, A3) and the NTS (B1, B2, B3) using confocal microscopy (x63 objective). Density of 5-HT positive fibers was calculated in the VLM (A4) and the NTS (B4). Data are expressed as the mean ± SEM (n=3 per group). *, P<0.05; ***, P<0.001, compared to controls and †††, P<0.001, compared to EPI/SDOC-.

#### Alteration in brainstem transcript level of TPH2 and SERT in EPI/SDOC+ rats

TPH2 transcript level was significantly increased in EPI/SDOC+ compared to both EPI/SDOC-rats and control rats. There was no statistical difference in transcript level of TPH2 between EPI/SDOC- and control rats (Figure 5A). SERT transcript level was significantly increased in EPI/SDOC+ compared to EPI/SDOC-rats, but not compared to control rats. In addition, SERT transcript level did not differ statistically between control and EPI/SDOC-rats (Figure 5B).

**Figure 5:**
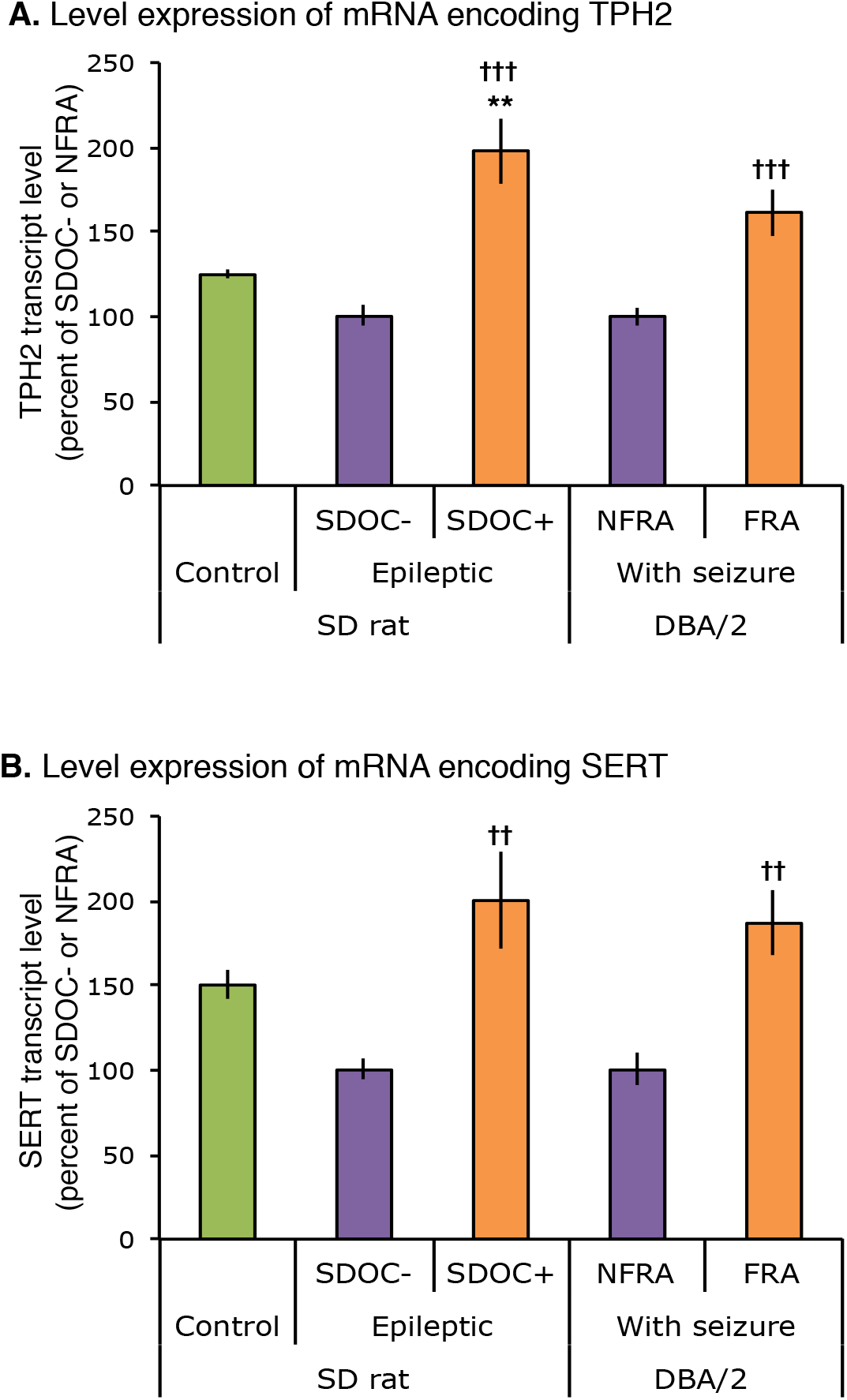
TPH2 and SERT mRNA transcript levels were measured in the brainstem of EPI rats and DBA/2 mice. Changes in brainstem levels of TPH2 (A) and SERT (B) transcripts were measured in EPI rats and DBA/2 mice at 8 weeks post-SE and 2-3 minutes after an audiogenic seizure (AGS), respectively. In contrast to EPI rats, DBA/2 mice had no history of seizure before AGS. Some mice developed a fatal respiratory arrest (DBA/2 with FRA) or a non-fatal respiratory arrest (DBA/2 with NFRA) after an AGS. TPH2 and SERT transcript levels in EPI/SDOC+ rats and in DBA/2 with FRA mice are expressed in percentage (mean ± SEM) of EPI/SDOC-rats and DBA/2 mice with NFRA, respectively (n=5 in each group). †: P<0.05, ††: P<0.01, †††: P<0.001, in comparison to EPI/SDOC-group for rat and DBA/2 with FRA group for mice. *, P<0.05; **, P<0.01, in comparison to control rats.

### Brainstem 5-HT system of DBA/2 mice prone to AGS

Previous studies demonstrated that almost 75% of DBA/2 mice developing AGS died from fatal respiratory arrest (FRA) following a tonic hindlimb extension [17]. Among the remaining mice, 15% developed non-fatal respiratory arrest (NFRA), meaning that they spontaneously recovered from post-AGS respiratory arrest. Brainstem 5-HT system of DBA/2 mice exhibiting FRA in response to AGS was shown to be altered in comparison with C57Black/6 mice that are resistant to AGS [10]. Thus, in experiment 4, we examined whether the alterations of TPH2 and SERT gene expression that we have observed between EPI/SDOC+ rats and EPI/SDOC-rats could be found between FRA and NFRA DBA/2 mice. Among the eleven male DBA/2 mice that underwent AGS with respiratory arrest, 64% of DBA/2 mice presented with FRA (n=7) and 36% with NFRA (n=4). We found that the levels of TPH2 and SERT transcripts were significantly elevated in FRA DBA/2 mice compared to NFRA DBA/2 mice (Figure 5A-B). It was striking that the level of TPH2 and SERT transcripts of FRA DBA/2 mice followed a trend that was similar to that of EPI/SDOC+ rats when both were compared to their respective controls (NFRA DBA/2 mice and EPI/SDOC-rats).

## Discussion

This study reports chronic alterations of respiratory system in epileptic (EPI) rats associated with 5-HT dysfunctions within the brainstem.

### Exploration of respiratory function in epilepsy models

To date, studies in animal models of seizures, aimed at understanding respiratory dysfunction in the context of epilepsy, were mostly performed during the time period surrounding seizures, i.e. during ictal and post-ictal periods [17–27]. Respiratory alterations have been detected in these abovementioned studies from short recording periods (2-30 min). In view of the respiratory disorders that can affect patients suffering from temporal lobe epilepsy (ref), this work carried out on a widely used preclinical model of this type of epilepsy (following status epilepticus induced by intraperitoneal administration of pilocarpine, Pilo-SE) had two major objectives.

The first was to study respiratory function in rats with a long history of epilepsy and at a distance from the period of brain remodeling that follows Pilo-SE-induced severe brain damages [28]. After a 10-week period of active epilepsy (12 weeks post-SE), we showed a decrease of breathing rhythm in epileptic (EPI) rats at rest. This result is different from that reported in a recent study showing no alteration in resting ventilation in rats 2-4 weeks after SE developed following intrahippocampal administration of pilocarpine [29]. However, beyond the fact that we did not use the same strain of rats (Wistar vs. Sprague-Dawley in our study) nor the same route of pilocarpine administration (intrahippocampally vs. intraperitoneally in our study), we agree with the authors of the above-mentioned study that the mechanisms leading to altered breathing may depend on the stage of epilepsy. In their study, the assessment was rather at an early, whereas it was clearly at a later stage of epilepsy in our study.

The second objective was to perform a long duration (24h) recording of the respiratory function in order to extend investigation beyond the period surrounding seizure. Using indirect respiratory thermochemistry, we achieved a 24-hour monitoring to determine the amount of oxygen consumed by the whole-body. Subsequently, we provide evidence of the presence of abnormal 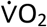 pattern in 30-50% of epileptic rats, which started with the installation of epilepsy (2-3 weeks post-SE). These EPI rats (EPI/SDOC+ group) presented with chronic 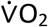 abnormalities characterized by low daily pattern among which intermittent and repetitive SDOC events were observed.

Until now, both preclinical and clinical studies investigated changes in the metabolic rate of O_2_ in epilepsy mostly at the brain level, reporting a decline in the activity of mitochondrial respiratory complexes in rats subjected to Pilo-SE and in patients with TLE [30, 31]. In our study, we measured the whole-body O_2_ intake which is mostly related to gas exchange during ventilatory process [32]. However, whole-body O_2_ intake can be also affected directly or indirectly by several factors including age, circadian rhythm, body-weight, bodytemperature, cardiac function and finally activity [16, 32–37]. None of these factors was found to be associated with abnormal pattern of 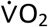 in our study. We then focused our attention on the ventilatory variables that may explain abnormal 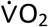 pattern in a subset of EPI rats.

### Respiratory system at rest in a subset EPI rats

The identification of several groups within a single disease is a common feature in human epilepsy, but rarely described in preclinical studies. One study has dichotomized two groups among rats subjected to kainic acid-induced SE according to the extent of brain damage and metabolic alterations [38]. In our study, EPI/SDOC+ subset has presented with a decrease of daily 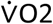 and a severe decrease of 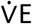 compared to EPI/SDOC-group.

During a 20-min monitoring period at rest, decreases of both 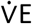 and 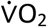 have already been described as an inherent dysfunction in a genetic model of audiogenic seizure in the WAR rat strain [26]. Because of technical limitations (24-hour 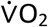 monitoring), we could not jointly assess 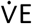 and 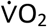 in our study, which prevented us from addressing the direct relationship that may exist between 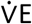 and 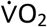. Nevertheless, we estimated for each rat the 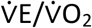 ratio as an indicator of ventilatory efficiency; this ratio was significantly higher in EPI/SDOC+ rats compared with EPI/SDOC- (+52%, P=0.007) and healthy control rats (+35%, P=0.04), and not statistically different between these two latter rat groups.

It is difficult at this stage to interpret these results, especially in the context of epilepsy. However, high 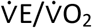 has already been described as an index of poor gas exchange during physical exercise of individual with heart failure disease [39]. Further exploration using blood oxygen measurement will be required to determine the direct relationship between 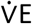 and 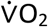 in epileptic rats following Pilo-SE. Another limitation of this study was the short duration of the ventilatory recording compared with the 24 hours of 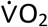 assessments. Significant fluctuations in 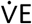 could have been observed, similar to those seen for 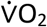, if the 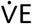 recordings had been longer. Further studies with devices allowing to concomitantly measure in free-moving rats both 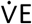 and 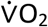 over sufficiently long periods of time will be needed to establish a direct link between 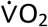 and 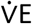 profiles and specifically during SDOC events.

Indeed, these events could represent a disruption of O_2_ uptake balance. But, to our knowledge, events similar to SDOCs have not yet been described and we can only interpret them in the light of what we know about a dyspneic situation. A first explanation for SDOCs could be the presence of ventilatory pauses; however, they would have to be extremely numerous and of long duration to induce such a collapse of 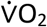. A second explanation could be a combination of several factors; for example, the absence of gasping, especially in hypoxic conditions, could lead to poor tissue oxygenation [40]. In EPI/SDOC+ rats, the 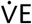 signal recording was too short in duration to determine whether the absence of gasping could be part of the ventilatory dysregulation.

### Ventilatory regulation in EPI rats

We observed an increased hypoxia ventilatory response (HVR) in EPI rats, in contrast to some studies that report no alteration in HVR in Pilo-SE rats [29, 41]. In addition, EPI/SDOC+ and EPI/SDOC-rats exhibited a sustained hypercapnia ventilatory response (HCVR) whereas previous studies have reported a diminished response in Pilo-SE-induced epilepsy, rapid amygdala kindling, and WAR rat models [26, 29, 42]. It is very difficult to compare our data with those in the literature, due to the fact that there are differences in models, timing of respiratory recordings, and severity of exposure to hypoxia and hypercapnia between our study and all others. Given that low HVR and HCVR are generally indicative of alterations in peripheral or central respiratory components, their maintenance in our study may indicate that if alterations occurred in the early phases, then they were compensated for or repaired in the late stages of epilepsy at which we performed our investigations.

### Brainstem 5-HT alterations

5-HT is mostly described as an excitatory drive toward respiratory network and subsequently the motor output [43]. Dysregulation of brainstem neuronal groups involved in the control of breathing, particularly those of the 5-HT system, has been found in patients with drugresistant epilepsy and has been associated with an increased risk of SUDEP [7]. We found that brainstem 5-HT innervation was impaired in EPI rats compared to control rats, with notable differences between EPI/SDOC+ and EPI/SDOC-rats. Indeed, EPI rats presented with an opposite 5-HT innervation pattern within the NTS. Since the impairment of respiratory function in the EPI/SDOC-rats seem to be moderate by opposition to EPI/SDOC+ rats, it is conceivable that, in these rats, the increase of 5-HT innervation in both the NTS and VLM may be part of protecting mechanisms of respiratory function. Such profile of 5-HT innervation was lacking in EPI/SDOC+ rats, but these rats presented with an increase of brainstem levels of both TPH2 and SERT transcripts. Such mechanisms in EPI/SDOC+ rats might support an increased concentration of 5-HT in neuron terminals by increasing 5-HT synthesis and re-uptake of intact 5-HT molecules, respectively, and thus may represent adaptive mechanisms to compensate, although not sufficiently, for the decreased brainstem 5-HT innervation. This conclusion is supported by results reported in conditional knock-out Lmx1b mice and WAR rats, which, as for EPI/SDOC+ rats, present similar respiratory alterations at rest and concomitant decrease of 5-HT drive [26, 43]. We also believe according to the difference of 5-HT innervation between VLM and NTS of the same EPI/SDOC+ rats that the decreasing of 5-HT could be structure specific and could be present in other primary regions of respiratory regulations.

We also compared brainstem 5-HT system of DBA/2 mice with FRA to their closest controls, DBA/2 mice with NFRA, contrasting with previous studies that compared DBA/2 mice with FRA to C57BL/6 mice resistant to AGS [10]. We found an increased level of mRNA coding for TPH2 and SERT in DBA/2 mice with FRA. This 5-HT profile in DBA/2 mice with FRA is intriguingly similar to that of EPI/SDOC+ rats. Altered 5-HT system is thus likely involved in respiratory dysfunctions in both acute and chronic seizure models.

In conclusion, Pilo-SE rat model, which shows both chronic respiratory alterations and spontaneous seizures, could be a relevant model to further explore neurobiological mechanisms underlying chronic alteration of respiratory regulation in epilepsy. Identifying the mechanisms that protect EPI/SDOC+ rats from mortality might also be an interesting approach to pave the way for new therapeutic strategies to prevent SUDEP.

## Methods

All animal procedures were in compliance with the guidelines of the European Union regulating animal experimentation (directive 2010-63), taken in the French law (decree 2013-118), and have been approved by the ethical committee of the Claude Bernard Lyon 1 University (protocol # BH-2008-11).

### Animals

Five week-old male Sprague-Dawley rats (Envigo, The Netherlands) were delivered to the animal facility of the Lyon 1 University (Villeurbanne, France) and housed in groups of 5 in transparent polycarbonate cages. Male DBA/2 mice of breeding colony of the LTSI laboratory were housed in groups of five. Rats and mice were maintained under a controlled 12-h/12-h light-dark cycle, in a temperature-controlled room (24-26 °C and 21-23 °C for rats and mice, respectively), with free access to food and water.

### Experimental design

Full description of methods have been described in details elsewhere [11, 28, 44–46]. Rats subjected to Pilo-SE are daily observed to detect spontaneous seizures until animals had developed epilepsy (EPI rats). Respiratory thermochemistry set-up was used to measure oxygen consumption 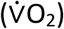 during 24 consecutive hours (daily 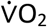) for each rat. Animals of this study were subjected to one of following experiments:

#### Experiment 1

To establish the time course of respiratory alterations occurring among PiloSE rats, 4 series of 10 rats were used in this experiment. In each series, Pilo-SE was induced in 8 rats and 2 rats were used as controls. Daily 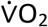 of each rat was measured once a week from the 1^st^ to the 8^th^ week post-SE.

#### Experiment 2

To further investigate cardio-respiratory functions in EPI rats during the chronic phase of epilepsy, Pilo-SE was triggered in 30 rats and 16 rats were used as controls. At the chronic period of epilepsy (12 weeks post-SE), the resting ventilation (normoxia, 21% O_2_) was measured using whole-body plethysmography. Between 13-14 weeks post-SE, a subset of EPI and control rats were used to evaluate ventilatory responses to hypoxia (HVR, FiO_2_= 9%) and to hypercapnia (HCVR, FiCO_2_= 5%), while others underwent surgery implantation of telemetric device to evaluate cardiac frequency and body temperature at 15 weeks post-SE. As 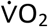 and ventilation are intermingled processes, to visualize global state of respiratory system in EPI rats, we had in-depth our data exploitation by characterizing ventilatory pattern according to 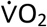 pattern. To note, daily 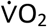 of EPI and control rats was measured short before plethysmography at 7-9 weeks post-SE.

#### Experiment 3

To determine whether integrity of the 5-HT system is altered in the brainstem of EPI rats presenting with respiratory alterations, Pilo-SE was induced in 20 rats and 8 rats were used as controls. Daily 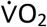 was evaluated between the 6^th^ and the 8^th^ week post-SE. Subsequently, animals were transcardially perfused and their brains were collected and processed either for histological procedures (5-HT immunodetection; n=3 in each group) or for gene expression analyses (SERT and TPH2 mRNA; n=5 in each group).

#### Experiment 4

To determine whether integrity of the 5-HT system is compromised in the brainstem of DBA/2 mice presenting with FRA, 11 DBA/2 mice underwent AGS followed by respiratory arrest. DBA/2 mouse brains were removed right after AGS for both FRA and nonfatal respiratory arrest (NFRA) DBA/2 mice. Mice with FRA were resuscitated and as for NFRA mice were injected with a lethal dose of pentobarbital and perfused with NaCl. Brainstem SERT and TPH2 gene expressions were measured and then compared between mice with either FRA or NFRA.

### *In vivo* procedures

#### Pilocarpine-induced SE

Epilepsy was triggered following Pilo-SE in 180-200 g male rats at the age of 7 weeks, as previously described [28, 45]. Scopolamine methylnitrate (1 mg×kg^−1^, s.c.; S-2250, Sigma-Aldrich) was administered 30 min prior to pilocarpine hydrochloride (350 mg×kg^−1^, i.p.; P-6503, Sigma-Aldrich). After an initial period of immobility, the onset of SE was characterized by repetitive tonic-clonic activity of the trunk and the limbs, occurring after repeated rearing with forelimb clonus and falling. Because SE event is a life-threating condition in rats, there is a need to stop it after 2 hours by a single injection of diazepam (Valium^®^, 10 mg×kg^−1^, i.p.; Roche). Rats were then hydrated with 2 mL of saline solution (0.9% NaCl; s.c.) and transferred to individual cages. After a one-week recovery period, all rats (control and treated rats) were housed in groups of 5 to avoid any stress that may result from social isolation.

#### Animal care after Pilo-SE

Control and treated rats were weighted daily during the first two weeks following Pilo-SE, and then every week until termination of the experiment. Daily abdominal massage was performed twice a day during the first week to activate intestinal motility, which is disrupted following Pilo-SE.

#### Detection of spontaneous recurrent seizures after Pilo-SE

To maintain as low as possible the level of anxiety in rats subjected to Pilo-SE, electro-encephalographic recordings (EEG) were not performed to detect the onset of epilepsy in our experiments. Indeed, conventional EEG set-up requires that rats are housed individually in a restrictive environment, which is known to enhance stress and anxiety and worsens epilepsy phenotype in Pilo-SE models [47]. In addition, as mentioned below in the “whole-body plethysmography” section, it is difficult to perform ventilatory recordings in animals with high anxiety level.

It is now clearly established that rats of the Sprague-Dawley strain from the Janvier Laboratories develop spontaneous recurrent seizures between 10 and 36 days after induction of Pilo-SE at 42 days of age [48, 49]. Because we were using Sprague-Dawley rats from Envigo Laboratories, we had to monitor the time of onset of spontaneous recurrent seizures (SRSs) in our experiments. To determine whether rats were in the chronic phase of epilepsy, they were manipulated daily between 09:00 and 11:00 a.m. to define the onset of handling-induced seizures (HIS), which was induced by restraining rats at the level of the chest with gentle pressure for 10 seconds [48]. HIS were used in our study to help experimenters to define the time to start detecting the first behavioural spontaneous recurrent seizures (SRSs). Thus, upon detection of 2 HIS, rats were observed by experimenters for 5 consecutive hours (between 01:00 and 06:00 p.m.) over the following days to detect the presence or absence of behavioural SRSs. Following Pilo-SE, rats are considered epileptic once they developed at least two SRSs during the periods of the daily observations. The severity of these seizures was scored from 3 to 5 on the Racine’s scale [50]. Seizure severity of less than 3 was difficult to detect visually in this study, which was conducted without daily video monitoring. Stage 3 corresponds to forelimb clonus, stage 4 to bilateral forelimb clonus with rearing, and stage 5 to rearing and falling or to tonic-clonic seizures.

During the 24 hour-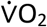 recordings, rats were video-monitored for 12h during the light period, making it possible to evaluate the number of seizures and their severity per session.

### Evaluation of respiratory and cardiac functions

#### Open respiratory thermochemistry

To detect respiratory alterations in epileptic rats, we used respiratory thermochemistry as an alternative approach to plethysmography, allowing respiratory function to be recorded for a long period of time on fed, unrestrained and unanesthetized animal. Respiratory thermochemistry mostly reflects gas exchanges and allows more specifically the measurement of the oxygen consumption 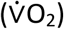 with no bias effect from animal’s body movements on the accuracy of respiratory acquisition.

Oxygen consumption 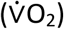 was measured with sample rate of 3 acquisitions per minute, under standard conditions (variation in temperature = 0°C; Ambient pressure = 760 mm Hg; dry air) using an open respiratory thermochemistry system, as previously described [51]. 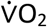 was continuously recorded during 24 consecutive hours on the same thermochemistry system; therefore, only one rat could be recorded each day in our experimental setting. Briefly, rats were placed individually in a thermostatic recording chamber (12 L) with steady airflow (4 L × min^−1^), under a controlled 12-h/12-h light-dark cycle (light from 06:00 a.m. to 06:00 p.m.), and were allowed free access to food and water. Ambient temperature inside the thermostatic chamber was monitored with copper-constantan thermocouples and was maintained at 24-26 °C using a thermostatic container. Airflow rate was measured using a Platon volumeter and converted to standard STPD (standard temperature and pressure, dry) values. Fractional O_2_ concentration was measured using a Servomex 1100 paramagnetic gas analyzer (Taylor Instrument Analytics ltd, Sussex, UK) and 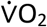 was calculated. The O_2_ analyzer was calibrated daily with pure nitrogen gas (0% O_2_) and room air (20.93% O_2_). Rats were weighted before and after recording then 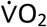 values were normalized to the average body weight (calculated from the weight measured before and after each recording session). Rats were video-monitored during the daylight period.

#### Whole-body plethysmography

Plethysmographic chamber allows collecting ventilatory variables from acquired breathing pattern signal. Ventilatory function was evaluated between 09:00 a.m. and 04:00 p.m. in unrestrained, unanesthetized rats under BTPS (body temperature and pressure, saturated) conditions using whole-body plethysmography (EMKA; technology). Calibration of the system was performed before each recording session by injecting a 1 mL air pulse inside the recording chamber. Briefly, each rat was individually placed in the recording chamber (600 mL) that is connected to a constant bias flow supply providing continuous inflow of fresh air (2.5 L × min^−1^). Minute volume (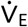; mL × min^−1^) was calculated as the product of tidal volume (V_T_; mL) and respiratory frequency (f_R_; breath x min^−1^). Total respiratory cycle duration (T_tot_) was calculated as the sum of the inspiratory time (T_I_; msec) and expiratory time (T_E_; msec). We also calculated for each animal the coefficient of variation of the respiratory frequency, which is an index of the variability of the respiratory pattern [52] and the frequency of spontaneous apnea, which is defined as a cessation of breathing lasting more than two respiratory cycles (>2 x T_tot_). Apnea number and duration were calculated. Apnea frequency was normalized to recording duration and expressed per min of breathing. Recordings were started after a 40-minute habituation period. Then, resting ventilation was monitored during 20 minutes under normoxic condition (21% O_2_). After recordings, rats were brought back to their home cage.

Ventilatory responses to hypoxia (FiO_2_= 9%) and to hypercapnia (FiCO_2_= 5%) were evaluated during two distinct recording sessions, which were performed a week apart from each other. After a 40-minute habituation period, plethysmographic recordings under basal condition (normoxia) were started 10 minutes before gas variations. Ventilatory responses to a 10minute exposure to hypoxia or hypercapnia were subsequently evaluated. All plethysmography recordings were made between 09:00 a.m. and 04:00 p.m.; because, beyond 04:00 p.m., EPI rats were more stressed in the plethysmography chamber, requiring longer time to reach a quiet state in this novel and restricted environment. At the end of each recording session, rats were brought back to their home cage. Repeated measurements are randomly performed for each rat.

#### Abdominal implantation of telemetry transmitters to evaluate cardiac function and bodytemperature

Rats were anesthetized with a mix of ketamine and xylazine (75 mg × kg^−1^ and 25 mg × kg^−1^, respectively; i.p.). A midline abdominal incision was made and a sterile telemetry transmitter (Data Sciences International, France) was placed inside the abdominal cavity. A pair of biocompatible wire electrodes originating from the transmitter were tunnelled subcutaneously and implanted to the left intercostal muscles and between the right shoulder blades. The transmitter was attached to the abdominal wall during wound closure with absorbable sutures. After surgery, animals received an intramuscular injection of penicillin (15,000 units) once a day for 3 consecutive days and were then returned to their home cage. Rats were monitored daily to control general health state. Two weeks after surgery, telemetry transmitters were magnetically activated and freely moving animals were placed in individual cages placed under platform-receiver units. Variation in cardiac frequency and body-temperature were recorded during 24 consecutive hours.

### Induction of audiogenic seizures in DBA/2 mice

AGS were induced as previously described [11, 17], with slight modifications. Briefly, at age of 23 days, mice were individually placed in a plastic cylinder (40 cm diameter, 20 cm height) and exposed to a broadband acoustic stimulus (intensity of 110 dB SPL with a peak frequency at 12 kHz) generated by a mechanical sonicator (Deltasonic model Delta 920). The stimulus was given for a maximum duration of 60 seconds or until the mouse exhibited AGS, characterized by an initial wild running and ending by a tonic hindlimb extension. All mice exhibited a tonic hindlimb extension leading to a respiratory arrest identified by pinnae relaxation and a complete cessation of chest movements. Respiratory arrest was followed by spontaneous recovery of the respiratory function within 5 seconds in mice identified as NFRA, or was followed by death for the other mice (FRA). Mouse brain was systematically removed within 2-3 min after acoustic stimulus was triggered.

### *Ex vivo* procedures

DBA/2 mice and rats were injected with a lethal dose of pentobarbital (250 mg × kg^−1^; i.p.) before being sacrificed. For gene expression analyses, animals were transcardially perfused (30 mL x min^−1^) for 2 min with ice-cold saline and their brain was rapidly removed. Then, the brainstem region was isolated, frozen in liquid nitrogen and stored at −80°C. For immunohistochemistry procedure, animals were transcardially perfused (30 mL x min^−1^) for 2 min with ice-cold saline and then for 10 min with ice-cold 4% paraformaldehyde in 0.2 M phosphate buffer (30 mL x min^−1^). After cryoprotection in a 25% sucrose solution, brains were frozen at −40°C in isopentane and stored at −80°C.

#### 5-HT immunohistochemistry

Coronal sections (40 μm thick) were cryostat-cut throughout the medulla oblongata and were processed free-floating for 5-HT immunostaining. Using the rat brain atlas, sections from each experimental group were selected at interaural coordinates −2.60 mm, and were all processed together for 5-HT immunofluorescent labeling. Selected sections were rinsed in 0.1M Phosphate Buffered Saline (PBS) and incubated in PBS containing 0.3% Triton X-100 and 3% normal donkey serum. Then, sections were incubated with a rabbit polyclonal antibody against 5-HT (Sigma; S5545; final dilution = 1:3,000). After few washes, sections were incubated with an AlexaFluor-488 conjugated donkey anti-rabbit antibody (Molecular Probes; A-21206; final dilution = 1:1,000). At the end of the immunostaining protocol, sections were mounted onto SuperFrost^®^Plus slides and cover slipped using Vectashield mounting medium (H-1500, Vector).

#### Semi-quantification of brainstem 5-HT immunostaining

Images (1152 x 1024 pixels) were captured with a HLX PL APO x63/0.70 objective and with the same conditions of photomultiplier gain, offset and pinhole aperture using a TCS SP2 confocal microscopy system (Leica), allowing the comparison of fluorescence intensity between sections. Image analysis was conducted on an average of 8 z-compressed images stacks (each image = 4 μm depth), which include the whole region of interest. Using the image J software (NIH), we first measured the average grey value within each 5-HT-positive neuronal cell body of caudal raphe nuclei (pallidus, magnus and obscurus), as an index of 5-HT neuronal concentration (A.U.). Using the same software, we also measured the density of 5-HT-immunopositive processes within the surface area of the NTS and the VLM, expressed as 5-HT-positive pixel/ pixel^2^.

#### Quantification of transcript level variation by RT-qPCR

Total RNA was extracted from brainstem samples using Tri-reagent LS (Molecular Research Center, Inc.). Contaminant genomic DNA was subsequently removed from the samples by treatment with Turbo DNA-*free*™ kit (Ambion). The messenger RNAs (mRNAs) contained in 1 μg of purified RNA extracts were reverse-transcribed using the reverse transcriptase RNase H minus (Promega) and oligo-d(T)15. To normalize the RT step, a synthetic external and non-homologous poly(A) standard RNA (SmRNA; Morales and Bezin, patent WO2004.092414) was added to the RT reaction mix (150,000 copies in each experimental sample), as previously described [28, 44–46]. PCR amplification of selected cDNAs, i.e. those complementary to the mRNA encoding rate-limiting enzyme of brainstem 5-HT synthesis (tryptophan hydroxylase: TPH2) and the 5-HT transporter (SERT), were performed using the Rotor-Gene Q system (Qiagen) and the QuantiTect SYBR Green PCR Kit (Qiagen). Sequences of the different primer pairs used for PCR amplification of mouse and rat TPH2 and SERT cDNAs are listed in Supplementary Table S1.

### Statistical analysis

The data for each experimental group were pooled and expressed as the mean ± SEM in the text and figures, unless otherwise specified. All statistics were performed using Sigma plot software 3.5 (Systat Software Inc., Point Richmond, California, USA). A Kolmogorov-Smirnov test was used to test if the data follows a normal distribution, then paired or unpaired t-test was performed for the two-group comparison. For multiple group comparisons, one-way ANOVA followed by post-hoc Fisher’s LSD test was used. Statistical significance was predefined as P<0.05.

## Abbreviations

5-HT: serotonin;
AGS: audiogenic seizures; EPI;
CIH: chronic intermittent hypoxia;
EEG: electro-encephalography; epileptic;
EPI/SDOC-: epileptic rat without sharp decrease in oxygen consumption;
EPI/SDOC+: epileptic rat with sharp decrease in oxygen consumption;
f_R_: respiratory frequency;
FRA: fatal respiratory arrest;
HCVR: hypercapnia ventilatory response;
HIS: handling-induced seizures;
HVR: hypoxia ventilatory response;
NFRA: non-fatal respiratory arrest;
GTCS: generalized tonic-clonic seizures;
NTS: nucleus tractus solitarius;
OSA: obstructive sleep apnea syndrome;
Pilo-SE: pilocarpine-induced *status epilepticus*;
SE: *status epilepticus*;
SERT: serotonin transporter;
SSRI: selective serotonin reuptake inhibitor;
SUDEP: sudden unexpected death in epilepsy;
T_E_: expiration time;
T_I_: inspiration time;
Tph2: tryptophan hydroxylase 2;
T_tot_: total respiratory cycle duration;
V_E_: minute volume;
VLM: ventrolateral medulla;
VO_2_: oxygen consumption;
V_T_: tidal volume

## Authorship

L.B and S.R. contributed equally to the article. L.B., S.R and P.R conceived and designed the experiments. A.M., J.R. and H.K performed respiratory thermochemistry experiments. H.K. performed plethysmography and telemetry experiments. B.M., G.D., L.B. and H.K., performed AGS in DBA/2 mice. H.K. and B.G. performed histochemistry and RT-qPCR experiments. L.B., S.R., M.O. and H.K. analyzed data. L.B., S.R. and H.K. wrote the paper.

## Acknowledgements

We thank for their assistance: Frédérique Cohen-Adad (scientific documentation; TIGER team, CRNL), and Denis Ressnikoff (confocal microscopy; Centre Commun de Quantimétrie of the Claude Bernard Lyon 1 University). Benoît Martin has received subventions from Eisai, UCB-Pharma and Zogenix.

## Conflicts of interest

None of the authors has any conflict of interest to disclose. We confirm that we have read the Journal’s position on issues involved in ethical publication and affirm that this report is consistent with those guidelines.

## Supplementary information

**Figure S1: Metabolic rate of oxygen at 1-2 weeks and 5-8 weeks post-SE.** The average oxygen consumption 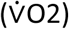 was calculated for the light and dark periods at 5-8 weeks post-SE (A) and then at 1-2 weeks post-SE (B) using the same experimental groups, retrospectively. Dichotomization of SDOC+ and SDOC-EPI rats was based on data collected at 5-8 weeks post-SE, which showed a significant decline in the average 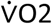 of EPI/SDOC+ rats during both the light and dark periods compared to EPI/SDOC- and control rats (A). There was no difference the three rat groups at 1-2 weeks post-SE, during which time no SDOCs were detected (B). Results are expressed in percent of control rats *, P<0.05; ***, p<0.001, compared to control rats. †††, P<0.001, compared to EPI/SDOC-rats.

**Figure S2: Ventilatory function in epileptic and control rats under normoxia**. (A) ventilation (V_E_), respiratory frequency (F_R_) and tidal volume (V_T_) of epileptic (n=16) and control (n=5) rats at 12 weeks post-*S£*. (B) Index of pattern stability includes coefficient of variation of ventilatory pattern and apnea number/duration. The results are expressed as the mean ± SEM. *, P<0.05; **, P<0.01, in comparison to control rats.

**Supplementary Table S1: Primer pairs used for qPCR amplification of TPH2 and SERT cDNAs of rat and mouse.**

## Key Points

- There are mild and severe patterns of respiratory dysfunctions among rats with epilepsy developed after pilocarpine induced *status epilepticus*.
- Severe respiratory dysfunctions were evidenced in 30-50% of rats with epilepsy.
- These dysfunctions consisted with intermittent episodes of sharp decrease in oxygen consumption (SDOC), slow breathing pattern and low metabolic rate of oxygen at rest.
- SDOCs were associated with decreased density of 5-HT-containing processes in the nucleus tractus solitarius.
- Rats with SDOCs had SERT and TPH2 transcript level alterations similar to DBA/2 mice subjected to AGS-induced fatal respiratory arrest.

## Notes

### Competing Interest Statement

The authors have declared no competing interest.

### Summary of Updates

This version is similar to the previous version, only author name was edited

## References

1. Lacuey, N., et al., The incidence and significance of periictal apnea in epileptic seizures. Epilepsia, 2018. 59(3): p. 573–582.

2. Bourdillon, P., et al., Letter to the Editor. Temporal lobe epilepsy: open or stereotactic surgery? J Neurosurg, 2019. 131(3): p. 989.

3. Ryvlin, P., et al., Incidence and mechanisms of cardiorespiratory arrests in epilepsy monitoring units (MORTEMUS): a retrospective study. Lancet Neurol, 2013. 12(10): p. 966–77.

4. Vilella, L., et al., Incidence, Recurrence, and Risk Factors for Peri-ictal Central Apnea and Sudden Unexpected Death in Epilepsy. Front Neurol, 2019. 10: p. 166.

5. Sivathamboo, S., et al., Sleep-disordered breathing in epilepsy: epidemiology, mechanisms, and treatment. Sleep, 2018. 41(4).

6. Somboon, T., M.M. Grigg-Damberger, and N. Foldvary-Schaefer, Epilepsy and Sleep-Related Breathing Disturbances. Chest, 2019. 156(1): p. 172–181.

7. Patodia, S., et al., The ventrolateral medulla and medullary raphe in sudden unexpected death in epilepsy. Brain, 2018. 141(6): p. 1719–1733.

8. Garcia, A.J., 3rd, et al., Chapter 3--networks within networks: the neuronal control of breathing. Prog Brain Res, 2011. 188: p. 31–50.

9. Faingold, C.L., et al., Differences in serotonin receptor expression in the brainstem may explain the differential ability of a serotonin agonist to block seizure-induced sudden death in DBA/2 vs. DBA/1 mice. Brain Res, 2011. 1418: p. 104–10.

10. Uteshev, V.V., et al., Abnormal serotonin receptor expression in DBA/2 mice associated with susceptibility to sudden death due to respiratory arrest. Epilepsy Res, 2010. 88(2-3): p. 183–8.

11. Martin, B., et al., Audiogenic seizure as a model of sudden death in epilepsy: A comparative study between four inbred mouse strains from early life to adulthood. Epilepsia, 2020.

12. Faingold, C.L., S. Tupal, and M. Randall, Prevention of seizure-induced sudden death in a chronic SUDEP model by semichronic administration of a selective serotonin reuptake inhibitor. Epilepsy Behav, 2011. 22(2): p. 186–90.

13. Tupal, S. and C.L. Faingold, Evidence supporting a role of serotonin in modulation of sudden death induced by seizures in DBA/2 mice. Epilepsia, 2006. 47(1): p. 21–6.

14. Curia, G., et al., The pilocarpine model of temporal lobe epilepsy. J Neurosci Methods, 2008. 172(2): p. 143–57.

15. Biagini, G., et al., Proepileptic influence of a focal vascular lesion affecting entorhinal cortex-CA3 connections after status epilepticus. J Neuropathol Exp Neurol, 2008. 67(7): p. 687–701.

16. Feher, J.J., Quantitative human physiology: an introduction. Second edition. ed. Academic Press series in biomedical engineering. 2017, Amsterdam; Boston: Elsevier/AP, Academic Press is an imprint of Elsevier. xiii, 968 pages.

17. Venit, E.L., B.D. Shepard, and T.N. Seyfried, Oxygenation prevents sudden death in seizure-prone mice. Epilepsia, 2004. 45(8): p. 993–6.

18. Aiba, I. and J.L. Noebels, Spreading depolarization in the brainstem mediates sudden cardiorespiratory arrest in mouse SUDEP models. Sci Transl Med, 2015. 7(282): p. 282ra46.

19. Jefferys, J.G.R., et al., Brainstem activity, apnea, and death during seizures induced by intrahippocampal kainic acid in anaesthetized rats. Epilepsia, 2019. 60(12): p. 2346–2358.

20. Faingold, C.L., M. Randall, and S. Tupal, DBA/1 mice exhibit chronic susceptibility to audiogenic seizures followed by sudden death associated with respiratory arrest. Epilepsy Behav, 2010. 17(4): p. 436–40.

21. Johnston, S.C., et al., Central apnea and acute cardiac ischemia in a sheep model of epileptic sudden death. Ann Neurol, 1997. 42(4): p. 588–94.

22. Kim, Y., et al., Severe peri-ictal respiratory dysfunction is common in Dravet syndrome. J Clin Invest, 2018. 128(3): p. 1141–1153.

23. Lertwittayanon, W., O. Devinsky, and P.L. Carlen, Cardiorespiratory depression from brainstem seizure activity in freely moving rats. Neurobiol Dis, 2019. 134: p. 104628.

24. Nakase, K., et al., Laryngospasm, central and obstructive apnea during seizures: Defining pathophysiology for sudden death in a rat model. Epilepsy Res, 2016. 128: p. 126–139.

25. St-John, W.M., et al., Changes in respiratory-modulated neural activities, consistent with obstructive and central apnea, during fictive seizures in an in situ anaesthetized rat preparation. Epilepsy Res, 2006. 70(2-3): p. 218–28.

26. Totola, L.T., et al., Impaired central respiratory chemoreflex in an experimental genetic model of epilepsy. J Physiol, 2017. 595(3): p. 983–999.

27. Villiere, S.M., et al., Seizure-associated central apnea in a rat model: Evidence for resetting the respiratory rhythm and activation of the diving reflex. Neurobiol Dis, 2017. 101: p. 8–15.

28. Sanchez, P.E., et al., Optimal neuroprotection by erythropoietin requires elevated expression of its receptor in neurons. Proc Natl Acad Sci U S A, 2009. 106(24): p. 9848–53.

29. Maia, O.A.C., et al., Pilocarpine-induced status epilepticus reduces chemosensory control of breathing. Brain Res Bull, 2020. 161: p. 98–105.

30. Kudin, A.P., et al., Seizure-dependent modulation of mitochondrial oxidative phosphorylation in rat hippocampus. Eur J Neurosci, 2002. 15(7): p. 1105–14.

31. Kunz, W.S., et al., Mitochondrial complex I deficiency in the epileptic focus of patients with temporal lobe epilepsy. Ann Neurol, 2000. 48(5): p. 766–73.

32. Gastinger, S., et al., A comparison between ventilation and heart rate as indicator of oxygen uptake during different intensities of exercise. J Sports Sci Med, 2010. 9(1): p. 110–8.

33. Gautier, H., Interactions among metabolic rate, hypoxia, and control of breathing. J Appl Physiol (1985), 1996. 81(2): p. 521–7.

34. Refinetti, R., Metabolic heat production, heat loss and the circadian rhythm of body temperature in the rat. Exp Physiol, 2003. 88(3): p. 423–9.

35. Refinetti, R., The circadian rhythm of body temperature. Front Biosci (Landmark Ed), 2010. 15: p. 564–94.

36. Seifert, E.L. and J.P. Mortola, Circadian pattern of ventilation during acute and chronic hypercapnia in conscious adult rats. Am J Physiol Regul Integr Comp Physiol, 2002. 282(1): p. R244–51.

37. Stephenson, R., et al., Circadian rhythms and sleep have additive effects on respiration in the rat. J Physiol, 2001. 536(Pt 1): p. 225–35.

38. Pearce, P.S., et al., Metabolic injury in a variable rat model of post-status epilepticus. Epilepsia, 2016. 57(12): p. 1978–1986.

39. Luks, A., R.W. Glenny, and H.T. Robertson, Introduction to cardiopulmonary exercise testing. 2013, New York: Springer. vii, 147 pages.

40. Poets, C.F., et al., Gasping and other cardiorespiratory patterns during sudden infant deaths. Pediatr Res, 1999. 45(3): p. 350–4.

41. Campos, R.R., F.R. Tolentino-Silva, and L.E. Mello, Respiratory pattern in a rat model of epilepsy. Epilepsia, 2003. 44(5): p. 712–7.

42. Totola, L.T., et al., Amygdala rapid kindling impairs breathing in response to chemoreflex activation. Brain Res, 2019. 1718: p. 159–168.

43. Hodges, M.R. and G.B. Richerson, Interaction between defects in ventilatory and thermoregulatory control in mice lacking 5-HT neurons. Respir Physiol Neurobiol, 2008. 164(3): p. 350–7.

44. Morales, A., et al., Unexpected expression of orexin-B in basal conditions and increased levels in the adult rat hippocampus during pilocarpine-induced epileptogenesis. Brain Res, 2006. 1109(1): p. 164–75.

45. Nadam, J., et al., Neuroprotective effects of erythropoietin in the rat hippocampus after pilocarpine-induced status epilepticus. Neurobiol Dis, 2007. 25(2): p. 412–26.

46. Navarro, F.P., et al., Brain heparanase expression is up-regulated during postnatal development and hypoxia-induced neovascularization in adult rats. J Neurochem, 2008. 105(1): p. 34–45.

47. Manouze, H., et al., Effects of Single Cage Housing on Stress, Cognitive, and Seizure Parameters in the Rat and Mouse Pilocarpine Models of Epilepsy. eNeuro, 2019. 6(4).

48. Dube, C., et al., Relationship between neuronal loss and interictal glucose metabolism during the chronic phase of the lithium-pilocarpine model of epilepsy in the immature and adult rat. Exp Neurol, 2001. 167(2): p. 227–41.

49. Leroy, C., et al., Temporal changes in mRNA expression of the brain nutrient transporters in the lithium-pilocarpine model of epilepsy in the immature and adult rat. Neurobiol Dis, 2011. 43(3): p. 588–97.

50. Racine, R.J., Modification of seizure activity by electrical stimulation. II. Motor seizure. Electroencephalogr Clin Neurophysiol, 1972. 32(3): p. 281–94.

51. Belouze, M., et al., Leanness of Lou/C rats does not require higher thermogenic capacity of brown adipose tissue. Physiol Behav, 2011. 104(5): p. 893–9.

52. Ogier, M., et al., Brain-derived neurotrophic factor expression and respiratory function improve after ampakine treatment in a mouse model of Rett syndrome. J Neurosci, 2007. 27(40): p. 10912–7.

