## Supplementary information for "Respiratory dysfunctions in two rodent models of chronic epilepsy and acute seizures and their link with brainstem serotonin system"

* co-last authors

Corresponding author:

Laurent Bezin, CRNL UMR5292 U1028, Epilepsy Institute IDEE, 59 boulevard Pinel, F-69500 Bron, France;; tel: +33 472 138 970.

### Results

#### Development of behavioral spontaneous recurrent seizures in rats after pilocarpine-induced *status epilepticus*

Briefly, in experiment 1, 32 rats were subjected to pilocarpine-induced *status epilepticus* (Pilo-SE); eight of them did not develop SE and four died during SE. By the end of the 2$^{nd}$ week post-SE, the 20 remaining rats developed spontaneous recurrent seizures (SRSs), indicating that they became epileptic (EPI). In experiment 2, 30 rats were subjected to Pilo-SE; 7 were excluded either because they did not develop SE (5/7) or because they died during SE (2/7). The remaining rats developed epilepsy (n=23 EPI rats) by the end of the 2$^{nd}$ week post-SE. In experiment 3, 20 rats were subjected to Pilo-SE; 2 of them did not develop SE and 2 others died during SE. The remaining rats (n=16) developed epilepsy by the end of the 2$^{nd}$ week post-SE.

#### Non-ventilatory variables related to $\dot{V}O_2$

In the 3 groups of rats (controls, EPI/SDOC+ and EPI/SDOC-), we evaluated the effect of three factors that have been referenced as affecting $\dot{V}O_2$ [1-7] : time ($t_0$ corresponds to the time of induction of SE in EPI rats), body weight and circadian rhythm. We also evaluated the effect of seizures on $\dot{V}O_2$.

***Time effect.*** It has been described in the main text. Time had no effect both on the frequency and the duration of SDOCs.

***Body-weight effect.*** Because $\dot{V}O_2$ of each rat is systematically reported to the body-weight, we analyzed body-weight variations in all groups of rats throughout the course of the experiments. Maximum body-weight loss in rats was observed three days after induction of SE, reaching 23 ± 2% of their initial weight (measured the day before SE). They then regained weight to return to control values from 49 days post-SE. It is to note that retrospective analysis of EPI rats showed no significant difference in body-weight variation between EPI/SDOC+ and EPI/DSOC- rats ($p=0.93$), indicating that the decrease in $\dot{V}O_2$ in EPI/SDOC+ rats cannot be related to a higher weight gain compared to EPI/SDOC-.

***Circadian rhythm effect.*** Since the daily $\dot{V}O_2$ pattern is known to be under the control of circadian rhythm [4-7], we compared $\dot{V}O_2$ between the light and the dark periods for each group of rats. At 1-2 weeks post-SE, the average $\dot{V}O_2$ increased during the dark period compared to the light period in all groups of rats (control rats: +21 ± 7 %, $p<0.001$; EPI/SDOC- rats: +28 ± 2 %, $p<0.001$; EPI/SDOC+ rats: +24 ± 7 %, $p=0.01$). At 5-8 weeks post-SE, such differences between the dark and the lights periods were still observed for the three groups of rats (control rats: +21 ± 2 %, $p=0.001$; EPI/SDOC- rats: +23 ± 3 %, $p<0.001$; EPI/SDOC+ rats: +17 ± 5 %, $p=0.006$). The regulation of circadian rhythm was statistically similar between groups. Regarding SDOCs occurrence, the severity of these events was not different between light and dark periods in EPI/SDOC+ rats.

***Seizure effect.*** To characterize the $\dot{V}O2$ pattern during seizure manifestation in EPI rats, we based our analysis on the concomitant 12h-daylight video-thermochemistry sessions. A total of 10 seizures occurred under video monitoring. These seizures have been exhibited by a total of 8 EPI rats (5 EPI/SDOC-, 3 EPI/SDOC+ rats). Similar to our previous visual observations during the first two weeks following SE, no significant difference was noticed for the number of seizures between EPI/SDOC+ and EPI/SDOC- rats ($p=0.52$). Both EPI/SDOC+ and EPI/SDOC- rats exhibited stage 4-5 seizure severity according to Racine's scale. The average duration of seizures was 77 ± 0.08 sec with no difference between EPI/SDOC+ and EPI/SDOC- rats ($p=0.67$). To determine $\dot{V}O_2$ variation during seizure in EPI/SDOC- and EPI/SDOC+ rats, the segments of $\dot{V}O_2$ before, during and after seizures were isolated for characterization. Short after seizure, $\dot{V}O_2$ increased to reach a plateau whose values were +61 ± 7 % higher than those measured during the 10 min prior to seizure occurrence. This increase was particularly sustained across time, with an average duration of 37 ± 5 min before returning to basal values. There was no correlation between seizure severity and the extent or duration of $\dot{V}O_2$ increase.

**Supplementary Table S1: Primer pairs used for qPCR amplification of TPH2 and SERT cDNAs of rat and mouse.**

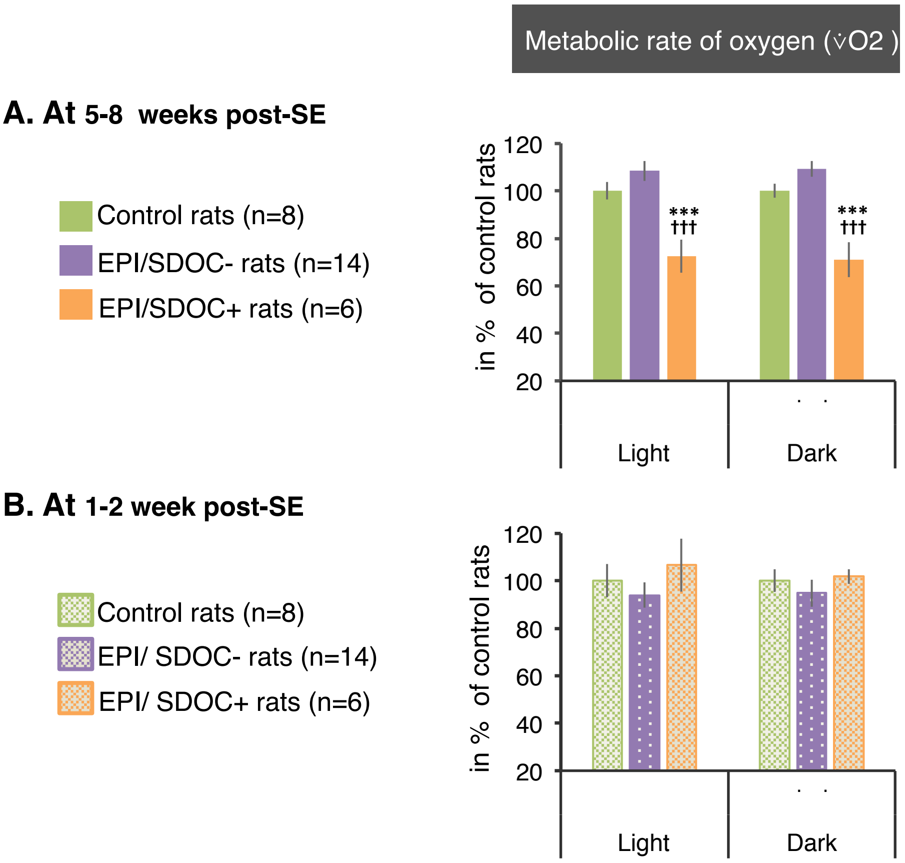


**Figure S1:** Metabolic rate of oxygen at 1-2 weeks and 5-8 weeks post-SE. The average oxygen consumption (v̇O2) was calculated for the light and dark periods at 5-8 weeks post-SE (A) and then at 1-2 weeks post-SE (B) using the same experimental groups, retrospectively. Dichotomization of SDOC+ and SDOC- EPI rats was based on data collected at 5-8 weeks post-SE, which showed a significant decline in the average v̇O2 of EPI/SDOC+ rats during both the light and dark periods compared to EPI/SDOC- and control rats (A). There was no difference the three rat groups at 1-2 weeks post-SE, during which time no SDOCs were detected (B). Results are expressed in percent of control rats *, $p<0.05$; ***, $p<0.001$, compared to controls. †††, $p<0.001$, compared to EPI/SDOC-.

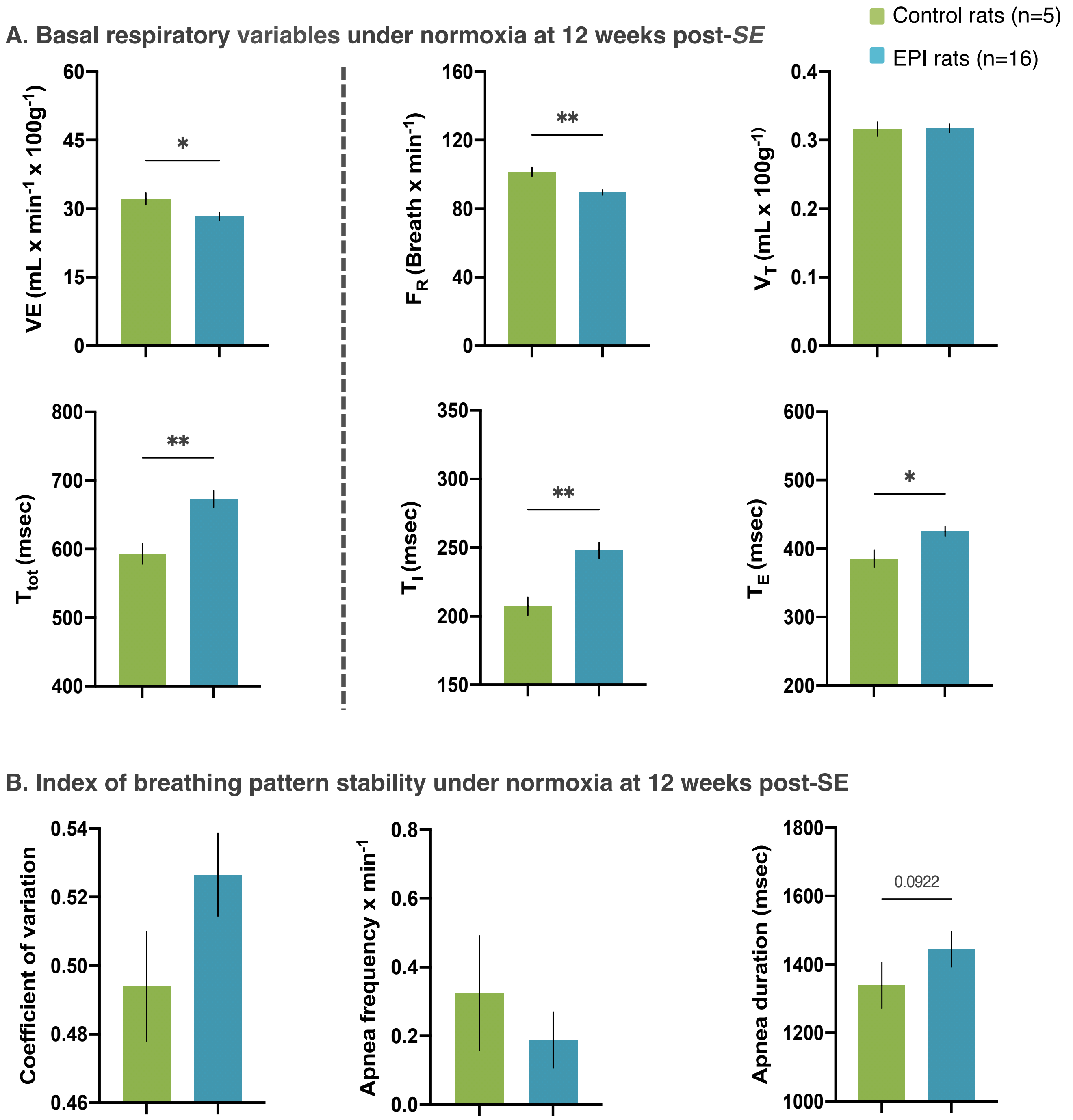


**Figure S2: Ventilatory function in epileptic and control rats under normoxia.** (A) ventilation ($V_E$), respiratory frequency ($F_R$) and tidal volume ($V_T$) of epileptic (n=16) and control (n=5) rats at 12 weeks post-*SE.* (B) Index of pattern stability includes coefficient of variation of ventilatory pattern and apnea number/duration. The results are expressed as the mean ± SEM. *, p<0.05; **, p<0.01, in comparison to control rats.

| Rat | Forward primer sequence (5´-> 3´) | Reverse primer sequence (5´-> 3´) |
|---|---|---|
| TPH2 (NM_173839.2) | TAC GGC ACC GAG CTT GAC | TGG CCA CAT CCA CAA AAT AC |
| SERT (NM_013034.3) | ATC ACC TGG ACG CTG CAT | TGG ATC TGC AGG ACA TGG |
| **Mouse** | **Forward primer sequence (5´-> 3´)** | **Reverse primer sequence (5´-> 3´)** |
| TPH2 (NM_173391.2) | GAG CTT GAT GCC GAC CAT | TGG CCA CAT CCA CAA AAT AC |
| SERT (NM_010484.2) | CAT ATG CTA CCA GAA TGG TGG A | AAG ATG GCC ATG ATG GTG TAA |

**Table S1: Primer pairs used for qPCR amplification of TPH2 and SERT cDNAs of rat and mouse.**
